# The autonomic effects of transcutaneous auricular nerve stimulation at different sites on the external auricle of the ear

**DOI:** 10.1101/2023.09.14.557755

**Authors:** Beatrice Bretherton, Aaron Murray, Susan Deuchars, Jim Deuchars

**Author notes:** Corresponding author: Beatrice Bretherton. These authors contributed equally (Beatrice Bretherton, Aaron Murray).

## Abstract

Transcutaneous auricular nerve stimulation (tANS) applied to specific parts of the external ear has positive health effects in both healthy volunteers and patient groups. However, due to differences between studies in ear stimulation sites and extent of effect on autonomic variables, it is challenging to determine what part of the external ear is the optimum site for electrode placement. This study investigated the autonomic effects of bilateral tANS at four different sites on the external auricle of the ear: tragus, cymba concha, helix and earlobe. tANS was performed using modified surface electrodes connected to a transcutaneous electrical nerve stimulation (TENS) machine. Participants (n = 24) each underwent four sessions where a 15 minute period of stimulation (pulse width: 200 μs; pulse frequency: 30 Hz; current: adjusted to sensory threshold) was applied bilaterally to either the tragus, cymba concha, helix or earlobe. Heart rate and blood pressure were continuously recorded during 10-minute baseline, 15-minute stimulation and 10-minute recovery periods. Heart rate variability (HRV) was derived. Results showed that regardless of site, stimulation significantly influenced measures of HRV. Baseline LF/HF ratio predicted change in LF/HF ratio during stimulation for each site and for all sites combined. Response (i.e. change in LF/HF ratio between baseline and stimulation) was closely linked with measures reflecting starting autonomic function. This demonstrates the importance of evaluating how autonomic function is modulated by tANS in individual participants (as well as the whole group). These findings have key implications for adopting a tailored approach when considering the therapeutic/clinical applications of tANS.

## INTRODUCTION

Transcutaneous auricular nerve stimulation (tANS) is a new and emerging technology that non-invasively stimulates specific parts of the external ear considered to contain sensory vagal nerves in order to promote health. This technology is well-tolerated by individuals, can be relatively inexpensive, is simple to administer and pain-free (1). There is increasing evidence that tANS can have positive effects on physiological and psychological function and may influence the autonomic nervous system. However, as tANS is relatively new, the present study aimed to elucidate if bilateral stimulation of different sites of the external ear could have similar effects on heart rate variability (HRV), considered a reflection of autonomic nervous system balance (2).

The external ear is innervated by at least three afferent nerves, the auricular branch of the vagus (ABVN), greater auricular nerve (GAN) and auriculotemporal nerve (ATN) (3). In this anatomical dissection of human cadaver ears, the authors described preferential innervation of different ear regions by the three nerves. According to the study, the cymba concha is wholly innervated by the ABVN, the tragus is equally innervated by the ABVN and GAN, with a minor contribution (9%) from ATN, the spine of helix is mainly innervated by ATN (with 9% GAN) while the lobule of auricle is wholly innervated by GAN. However there have been questions about inconsistencies in this report (4) particularly regarding innervation pattern of the ABVN.

Such questions on anatomical innervation are relevant when considering data from functional studies using the two stimulation sites that were originally thought to be significantly innervated by the ABVN, the tragus and cymba concha. tANS delivered bilaterally to the tragus of the external ear can modulate measures of HRV (5–9) and further investigation suggested that this is due to both increases in parasympathetic activity (indicated by increases in baroreflex sensitivity (6)) and decreases in muscle sympathetic activity (5). Furthermore, tANS delivered by stimulation of tragus is associated with significant health benefits in cardiovascular conditions, such as paroxysmal atrial fibrillation (10,11) and stroke (12). Similarly, cymba concha stimulation improves stroke recovery (13) and hypertension (14). However, there are inconsistencies in the literature regarding effectiveness of cymba concha stimulation on autonomic variables, perhaps dependent on differences in the study design. Importantly, as recently advocated (15), inconsistencies between studies could be due to a number of factors, such as unilateral vs. bilateral stimulation, and different stimulation parameters (e.g. pulse width, pulse frequency, current) and perhaps in part influenced by different devices used to generate the stimulation. For example, the effectiveness of right cymba concha stimulation on HRV was dependent on duration of stimulation and gender of subjects (16). However, no differences in measures of cardiac vagal activity were reported between cymba concha (considered “active”) and earlobe (considered “sham”) stimulation (15,17–19). This lack of difference between “active” and “sham” could be partly because the earlobe is innervated by the GAN, which feeds into cervical plexus and this may also influence autonomic variables, as suggested by Rangon (20,21). This would also be consistent with studies in rat where cervical sensory nerves mediated the decreased sympathetic activity elicited by tragus stimulation (22). Towards addressing such issues, unilateral stimulation at different sites of the ear increased HRV in a charge-dependent fashion, with stimulation at the right ear having a greater effect than the left and strongest effects from stimulation of the cymba concha, fossa triangularis and the inner tragus (23). Thus, tANS stimulation, regardless of site, may influence autonomic activity, not just through activation of the ABVN but also through the other sensory nerves that can influence autonomic circuits at different points in central pathways.

Considering the uncertainty surrounding sensory innervation of the external auricle and differences in responses observed with stimulation sites, this study aimed to investigate how HRV measures changed with bilateral stimulation administered to four different ear sites suggested by anatomical studies (3) to be differentially innervated by the 3 afferent nerves:

1. Cymba concha; a frequent target for tANS.
2. Tragus; another frequent target of tANS.
3. Earlobe; frequently used as a sham control.
4. Helix; considered to be predominantly innervated by the ATN and therefore potentially another sham control.

Since not all participants respond to tANS at the tragus with increases in measures of HRV (5,9), we also examined whether it was possible to identify potential responders based on initial HRV measures. Lastly, in response to Borges et al.’s call (15) for exploring subjective experiences of tANS, we undertook an exploratory investigation into how the stimulation at the four ear sites was perceived by participants.

It was hypothesised that:

- Stimulation to tragus, cymba concha and earlobe would be associated with changes in measures of HRV, compared to helix stimulation.
- Baseline measures of HRV would influence magnitude and direction of response to tANS.

## MATERIALS AND METHODS

### Ethical approval

University of Leeds Ethical Review Committee approved the project (Reference: BIOSCI 16-009), which conformed to Declaration of Helsinki standards. Participants provided informed written consent. Anonymised data were stored securely according to the UK Data Protection Act (1998). Participants were free to withdraw from the study at any point.

### Participants

Volunteers with a history of hypertension, cardiovascular disease, epilepsy or diabetes mellitus were excluded. Participants were requested to have no caffeine, alcohol, nicotine or strenuous exercise for at least 12 hours prior to visits. Hormone replacement therapy (HRT) was not being taken by any participants. To assess participant numbers, a Wilcoxon signed-rank test sample size calculation was performed. To detect a difference in measurements of tANS treatment with an effect size of 0.5, power of 60% and a significance level of 5%, 23 participants were required. To account for potential attrition, 24 healthy volunteers were recruited (aged ≥ 18 years; mean age: 33.09 ± 2.18 years, gender equally split).

### Procedure

Studies were conducted between the hours of 08.00 and 10.00 in a quiet, temperature controlled (21 ± 2°C) room at the University of Leeds, with participants reclined semi-supine on a couch.

### Transcutaneous auricular stimulation

A TENS machine (V-TENS Plus, Body Clock Health Care Ltd, United Kingdom) was used to deliver stimulus to left and right ears simultaneously, via customised auricular electrode clips (Auricular Clips, Body Clock Health Care Ltd, UK attached to the tragus, helix and earlobe). For the cymba concha, two carbon fibre electrodes were applied separately to the skin, positioned approximately 5 millimetres apart (see Figure 1) and held securely in place with a non-conductive silicone putty ear plug moulded to the shape of the participant’s cymba concha. This putty was then held in position using a lightweight headset to ensure stability and good contact between the electrodes and the skin. This ensured that testing was not interrupted due to potential movement of the cymba concha stimulation set-up. Participants wore the electrode clips throughout all the proceedings (baseline, stimulation, recovery). Stimulation parameters were detailed in prior work (5,9): 15 minutes continuous, pulse width 200 μs and pulse frequency 30 Hz, amplitude was adjusted to the level of sensory threshold (usually 2-4 mA). Participants were asked to report the sensations described during the initial titration of the stimulus at each site. When the stimulus was clearly detected by the participant, the current was turned down until the stimulus was borderline perceptible. Participants were also asked whether the sensation was localised to the area immediately in contact with the electrode or if the stimulus could be perceived elsewhere.

**Figure 1:**
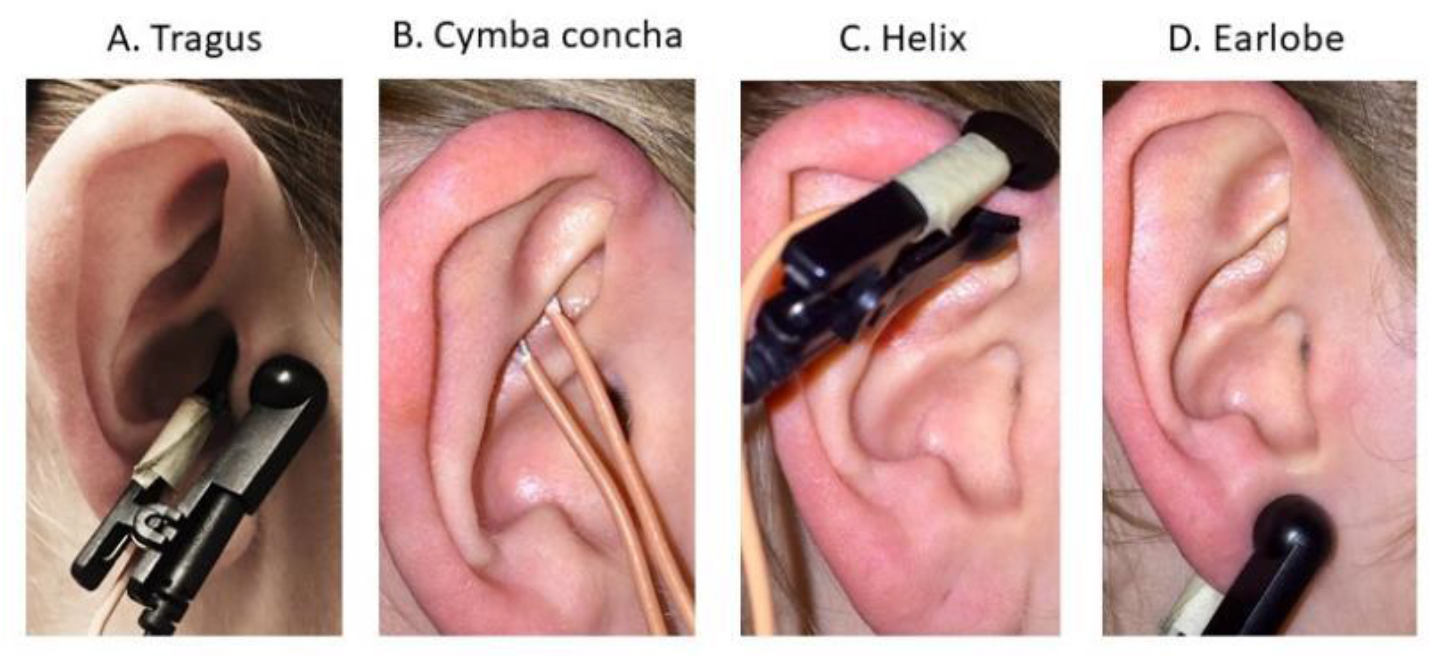
Electrode placement for the tragus (A), cymba concha (B), helix (C) and earlobe (D).

### Procedure

Participants attended on four occasions, one week apart, to compare the autonomic effects of transcutaneous auricular stimulation at four sites on the external ear (one site per visit): tragus, helix, cymba concha and earlobe. The order of the visits was randomized between participants. At the beginning of the first visit, participants completed a basic health questionnaire, physical activity level was assessed using the Godin Leisure Time Exercise Questionnaire (24) and height and weight obtained. Physiological equipment to continuously record heart rate, blood pressure and respiration was attached. Consistent with prior research (5,9), three sets of recordings obtained during each visit included 10 minute baseline, 15 minute stimulation and 10 minute recovery. Figure 2 summarises the procedure employed in the study.

**Figure 2:**
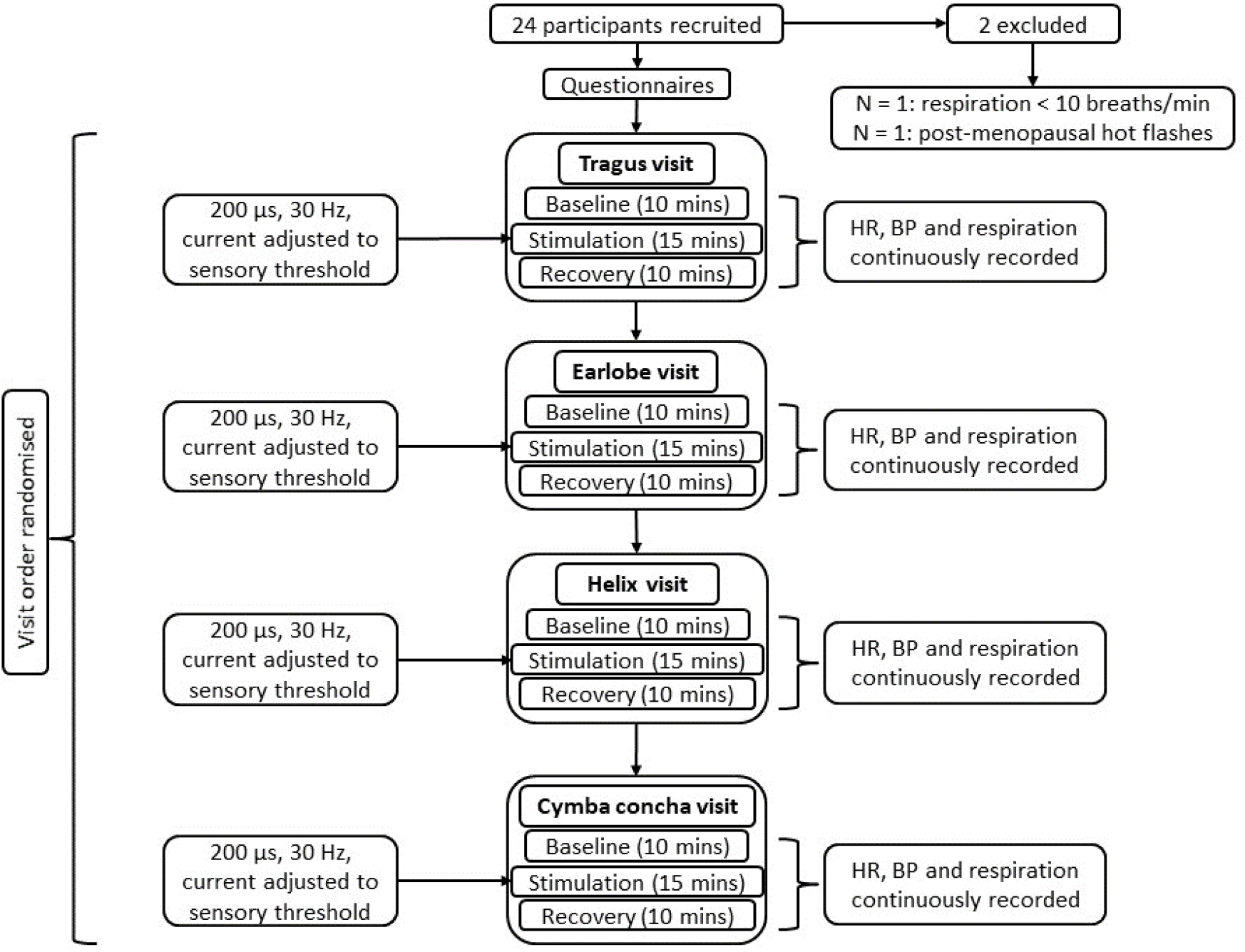
Summary of study procedure.

### Measurements

Continuous recordings were made of heart rate (HR), non-invasive blood pressure (BP) and respiration during baseline, stimulation and recovery. Non-invasive BP was recorded for the primary purpose of deriving baroreceptor reflex sensitivity (BRS). As frequency-domain HRV can be modulated by respiratory changes (25), the primary purpose of measuring respiration was to ensure that respiration rates did not drop to fewer than 10 breaths/minute and therefore was not analysed in depth. The final five minutes of each recording was used to derive frequency-domain, time-domain and non-linear HRV and cardiac BRS offline in LabChart 8 (ADInstruments). Change (Δ) between baseline and stimulation was calculated in all four visits.

### HRV variables Time-domain HRV

LabChart 8 was used to derive the following time domain measures: mean RR interval, Δ RR (difference between the longest and shortest RR intervals), SDRR (standard deviation of all RR intervals), RMSSD (the square root of the squared of differences between adjacent RR intervals) and pRR50 (percentage of number of pairs of adjacent RR intervals differing by more than 50 ms). SDRR reflects global autonomic regulation and is an estimate of all HRV (2). RMSSD and pRR50, being mainly related to beat-to-beat variations, are considered to reflect parasympathetic output (2).

### Frequency-domain HRV

Frequency-domain HRV parameters included: the low frequency (LF) component, detected at 0.04-0.15 Hz; the high frequency (HF) component, detected at 0.15-0.40 Hz and total power (0.04-0.40 Hz). Currently, there is controversy concerning what physiological system LF power may reflect. For instance, it has previously been proposed that LF power provides a means to indirectly evaluate the ability of the autonomic nervous system to modulate both sympathetic and parasympathetic outflow (26). However, understanding that is more recent proposes that LF power may reflect baroreflex modulation of autonomic outflow rather than cardiac sympathetic innervation (27–29). The HF component has been associated with parasympathetic modulation of heart rate (26). The ratio of LF to HF power (LF/HF) was also calculated.

### Non-linear HRV

Non-linear HRV has been advocated as a complementary indicator of HRV (29). This entails generating a Poincaré plot, in which consecutive RR intervals are plotted against the following interval to provide a two-dimensional graphic representation of the correlation between consecutive RR intervals. An ellipse fitted to the shape formed by the plot with the centre determined by the average RR intervals is used to provide quantitative measures SD1 and SD2. SD1 measures the standard deviation of the distances of the points to the diagonal y=x and SD2 measures the standard deviation of the distances of points to the line y=-x + RRm, where RRm is the average of RR intervals. SD1 is thought to represent parasympathetic activity as it is an index of instantaneous recording of beat-to-beat variability, while SD2 has been associated with the variability of long-term variability (30). SD1 and SD2 were normalised relative to heart rate by computing: (SD1 or SD2/RR interval)*1000. SD2/SD1 was also derived and represented the ratio between short- and long-term variations in RR intervals. The area of the ellipse, S, representing total variability in RR intervals was ascertained by calculating π*SD1*SD2.

### Cardiac baroreflex sensitivity (BRS)

The sequence method was applied to assess cardiac BRS (31). A sequential rise in both SBP (≥ 1 mmHg) and RR interval (≥ 2 ms) for three or more consecutive cardiac cycles described ‘Up’ sequences. In contrast, three or more cardiac cycles for which there was a sequential fall in SBP and RR interval were ‘Down’.

Plotting the RR interval against SBP for each sequence (R^2^ ≥ 0.85 acceptance level) and the average slope values for the ‘up’ and ‘down’ sequences were combined to get an average cardiac baroreflex slope. We accepted values of cardiac BRS when the number of sequences was ≥ 3 for both up and down sequences.

### Statistical Analysis

SPSS (version 24) was used for statistical analysis, Shapiro-Wilk for testing of normality. Two-tailed statistical tests with an alpha level of 0.05 were used in all instances. Unless otherwise stated, data are presented as group mean ± 1 standard error of the mean (SEM).

To explore differences between the ear stimulation sites (tragus, cymba concha, helix, earlobe) and the three conditions (baseline, stimulation, recovery), two-way repeated measure ANOVAs were performed. Mixed mode ANOVAs also explored differences between response type (responder, non-responder) and the three conditions (baseline, stimulation, recovery). The Greenhouse Geisser correction was used when data did not meet sphericity assumptions. To correct for multiple comparisons following statistically significant effects, the Bonferroni correction was applied to all post-hoc tests. Linear regression modelling explored the extent to which baseline autonomic function significantly predicted response to stimulation for all and individual ear sites.

## RESULTS

Twenty-four participants with no previous medical history of hypertension, cardiovascular disease, diabetes or epilepsy were enrolled. One male participant (age = 28 years) was excluded from the study due to a consistently slow respiration rate (< 8 breaths per minute) and non-compliance with a breathing metronome following coaching. One female participant (age = 62 years) was excluded from the study due to post-menopausal hot flashes (2-3 per visit, mean duration = 2 minutes) which generated frequent ECG signal artefacts. The baseline characteristics of the remaining 22 participants (n = 11 female and n = 11 male) included in the study are presented in Table 1.

**Table 1:** Summary of characteristics of the final study sample. Data presented as mean ± 1 SEM unless otherwise stated.

|  |  |
| --- | --- |
| Final sample size (n) | 22 |
| Gender (n males) | 11 |
| Age (yrs.) | 33.09 ± 2.18 |
| BMI (kg/m<sup>2</sup>) | 24.66 ± 0.84 |

### tANS significantly influenced HRV

Comparing stimulation to the baseline and recovery conditions for all stimulation sites together revealed that time-domain measures of HRV changed (Table 2). For instance, there was a significant main effect of condition on RR interval (F(1.16,24.37) = 33.66, p < 0.001, η _p_^2^ = 0.616), Δ RR (F(2,42) = 10.96, p < 0.001, η_p_^2^ = 0.343), SDRR (F(1.29,27.07) = 15.17, p < 0.001, η_p_^2^ = 0.419), RMSSD (F(1.16,24.29) = 12.11, p = 0.001, η_p_^2^ = 0.366) and pRR50 (F(1.44,30.26) = 11.36, p = 0.001, η_p_^2^ = 0.351).

**Table 2:** Significant main effects of condition emerged for RR interval (p < 0.001), Δ RR (p < 0.001), SDRR (p < 0.001), RMSSD (p = 0.001) and pRR50 (p = 0.001). Pairwise comparisons revealed significant differences between baseline, stimulation and recovery. * = significantly different to baseline; # = significantly different to stimulation. Data presented as mean ± 1 SEM.

|  | Baseline | Stimulation | Recovery |
| --- | --- | --- | --- |
| RR interval (ms) | 890 ± 30 | * 946 ± 28 | * # 961 ± 28 |
| $\Delta$ RR (ms) | 322 ± 35 | * 363 ± 40 | * 381 ± 37 |
| SDRR (ms) | 59.92 ± 7.18 | * 66.79 ± 7.91 | * # 72.00 ± 8.12 |
| RMSSD (ms) | 49.14 ± 7.93 | * 58.89 ± 8.81 | * 60.77 ± 8.66 |
| pRR50 (ms) | 24.78 ± 4.54 | * 31.38 ± 4.90 | * 31.72 ± 4.74 |

Further analysis revealed that RR interval, Δ RR, SDRR, RMSSD and pRR50 significantly increased between baseline and stimulation (p < 0.001, p = 0.023, p = 0.023, p = 0.008 and p = 0.001 respectively) and between baseline and recovery (p < 0.001, p = 0.002, p = 0.001, p = 0.004 and p = 0.009 respectively). RR interval and SDRR were also significantly higher during recovery compared to stimulation (p = 0.005 and p = 0.001 respectively).

Non-linear indices of HRV also varied between the conditions (Table 3). For instance, there was a significant main effect of condition on S (F(1.07,22.46) = 7.49, p = 0.011, η_p_^2^ = 0.263), SD1 (F(1.16,24.29) = 12.12, p = 0.001, η_p_^2^ = 0.366), nSD1 (F(1.20,25.09) = 9.87, p = 0.003, η_p_^2^ = 0.320), SD2 (F(1.39,29.21) = 14.72, p < 0.001, η_p_^2^ = 0.412) and nSD2 (F(2,42) = 10.76, p < 0.001, η_p_^2^ = 0.339).

**Table 3:** Significant main effects of condition emerged for S (p = 0.011), SD1 (p = 0.001), nSD1 (p = 0.003), SD2 (p < 0.001) and nSD2 (p < 0.001). Pairwise comparisons revealed significant differences between the three conditions. * = significantly different to baseline; # = significantly different to stimulation. Data presented as mean ± 1 SEM.

|  | Baseline | Stimulation | Recovery |
| --- | --- | --- | --- |
| **S (ms<sup>2</sup>)** | 11579 ± 3509 | 15123 ± 4273 | * # 16656 ± 4480 |
| **SD1 (ms)** | 34.80 ± 5.62 | * 41.72 ± 6.24 | * 43.04 ± 6.13 |
| **nSD1** | 36.87 ± 5.44 | * 42.54 ± 5.80 | * 43.46 ± 5.66 |
| **SD2 (ms)** | 75.21 ± 8.57 | * 84.24 ± 9.42 | * # 91.74 ± 9.86 |
| **nSD2** | 81.40 ± 7.88 | 87.14 ± 8.53 | * # 93.81 ± 8.95 |

Further analysis revealed that SD1, nSD1 and SD2 significantly increased between baseline and stimulation (p = 0.008, p = 0.014 and p = 0.040 respectively), and between baseline and recovery (p = 0.004, p = 0.011 and p = 0.001 respectively). S, SD2 and nSD2 were also significantly greater during recovery than stimulation (p = 0.002, p = 0.001 and p = 0.002 respectively) with S and nSD2 also being higher during recovery than baseline (p = 0.021 and p = 0.007 respectively).

### Similar effects on HRV were observed for all stimulation sites

Differences between the ear stimulation sites (tragus, earlobe, helix and cymba concha) did not reach statistical significance (main effect of ear stimulation site, p > 0.05). There was also little influence of stimulation at the different ear sites (condition*ear stimulation site interaction effects, p > 0.05). For a summary of all main effect and interaction effect outcomes, please see supplementary Table 1.

### Baseline LF/HF predicted change in LF/HF during tANS

Using frequency-domain HRV measures, there were no overall effects when comparing between sites and conditions. Therefore, since baseline LF/HF ratio has previously been shown to be a predictor of response (albeit for stimulation delivered to the tragus (5,9)), linear regression modelling was performed on the final sample for all ear stimulation sites. Indeed, findings revealed that for all participants regardless of ear stimulation site, baseline LF/HF ratio significantly predicted change in LF/HF ratio (R^2^ = 0.543, p < 0.001, Figure 3).

**Figure 3:**
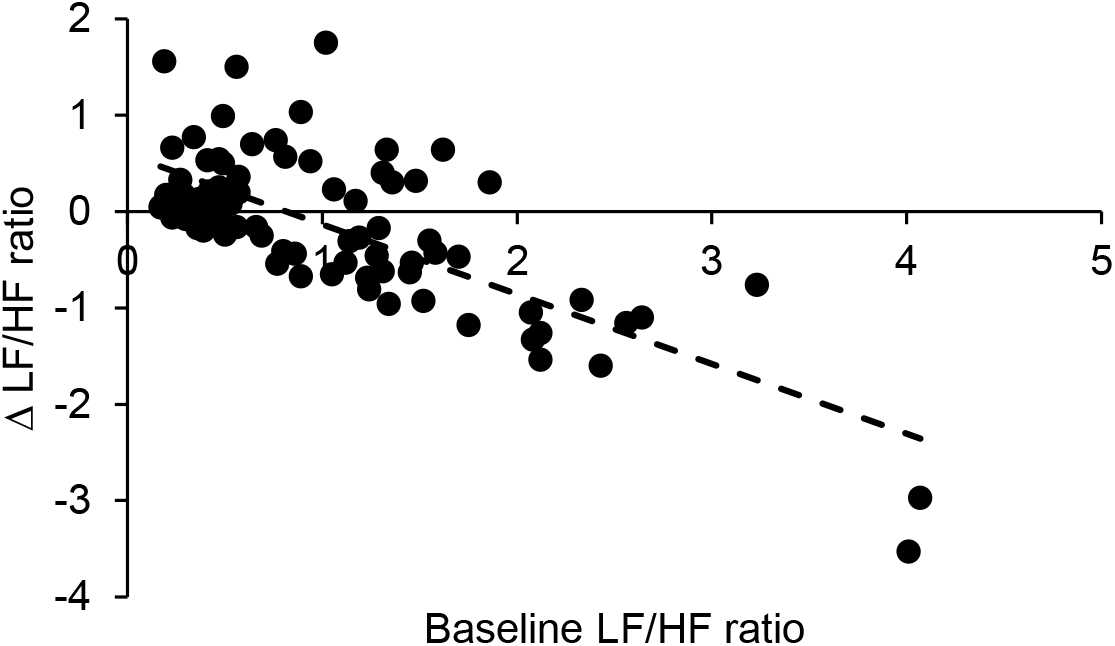
Regardless of ear site, baseline LF/HF ratio significantly predicted change in LF/HF ratio during stimulation.

Intriguingly, similar patterns emerged for the individual ear stimulation sites (including other HRV measures, see Supplementary Table 2 for a summary). For instance, baseline LF/HF was a significant predictor of change in LF/HF with stimulation to the tragus (R^2^ = 0.336, p = 0.005), cymba concha (R^2^= 0.793, p < 0.001), helix (R^2^ = 0.349, p = 0.004) and earlobe (R^2^ = 0.388, p = 0.002). Consistent with prior work (5,9), higher baseline LF/HF ratios were associated with greater decreases during stimulation at all sites (Figure 4).

**Figure 4:**
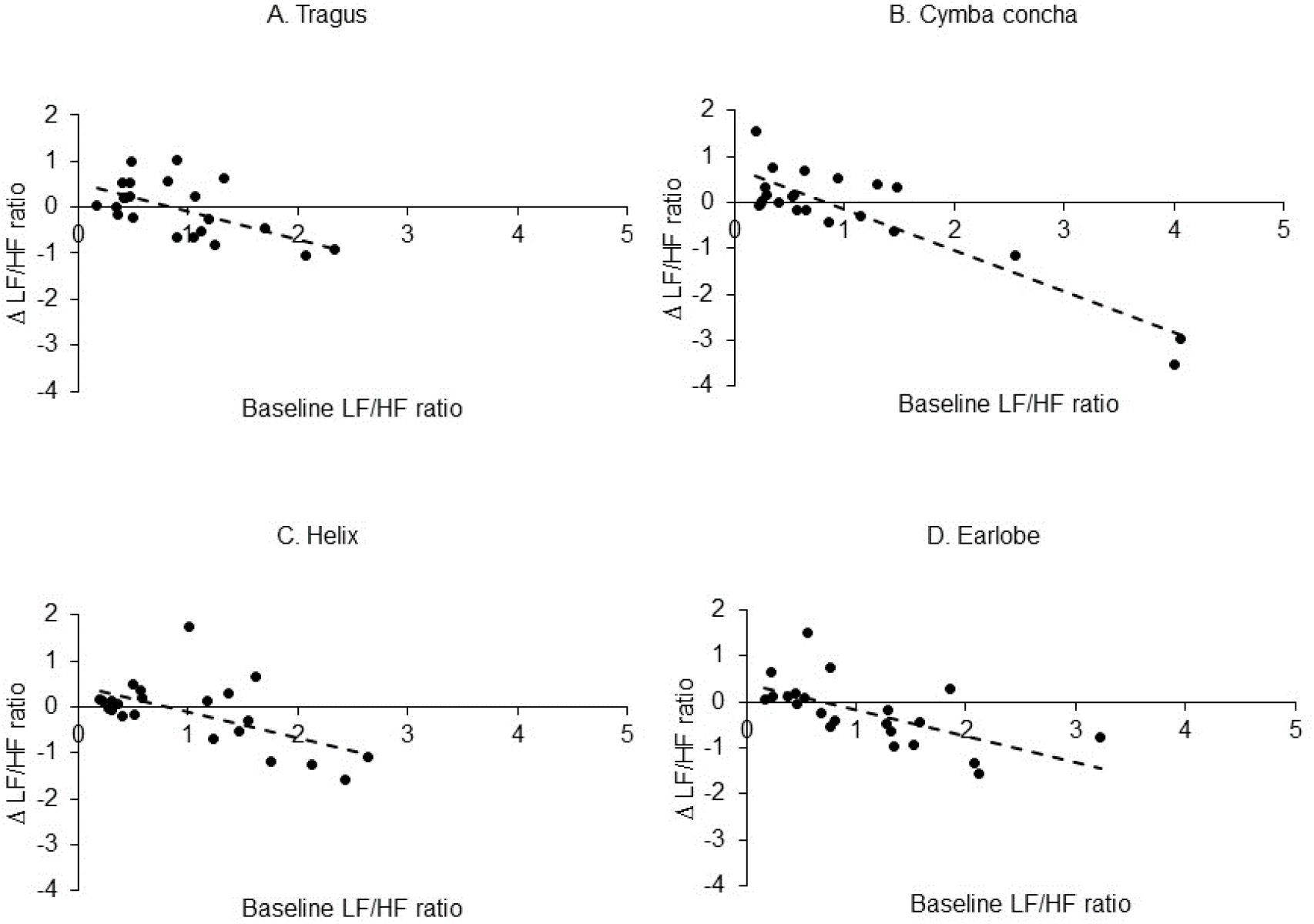
Baseline LF/HF ratio significantly predicted change in LF/HF ratio during stimulation at the tragus (A), cymba concha (B), helix (C) and earlobe (D).

### Responders had higher baseline LF/HF ratios than non-responders

Based on the definition of a response to tANS at the tragus (5,9), participants who showed a decrease ≥ 20% in LF/HF ratio (between baseline and stimulation) were classified as responders whilst those who showed increases or a < 20% decrease in LF/HF ratio were classified as non-responders.

The response rate for the four ear stimulation sites was as follows: 45.45% of participants responded to tragus stimulation; 40.91% for cymba concha stimulation; 40.91% for helix stimulation and 50% for earlobe stimulation.

We then explored how LF/HF ratio differed between response types and conditions. Indeed, mixed mode ANOVAs revealed significant interaction effects between response type and condition for stimulation delivered to the tragus (F(2,40) = 11.13, p < 0.011, η_p_^2^ = 0.358), cymba concha (F(2,40) = 11.02, p < 0.011, η_p_^2^ = 0.355), helix (F(2,40) = 9.49, p < 0.011, η_p_^2^ = 0.322) and earlobe (F(2,40) = 9.89, p < 0.011, η_p_^2^ = 0.331, Figure 5).

**Figure 5:**
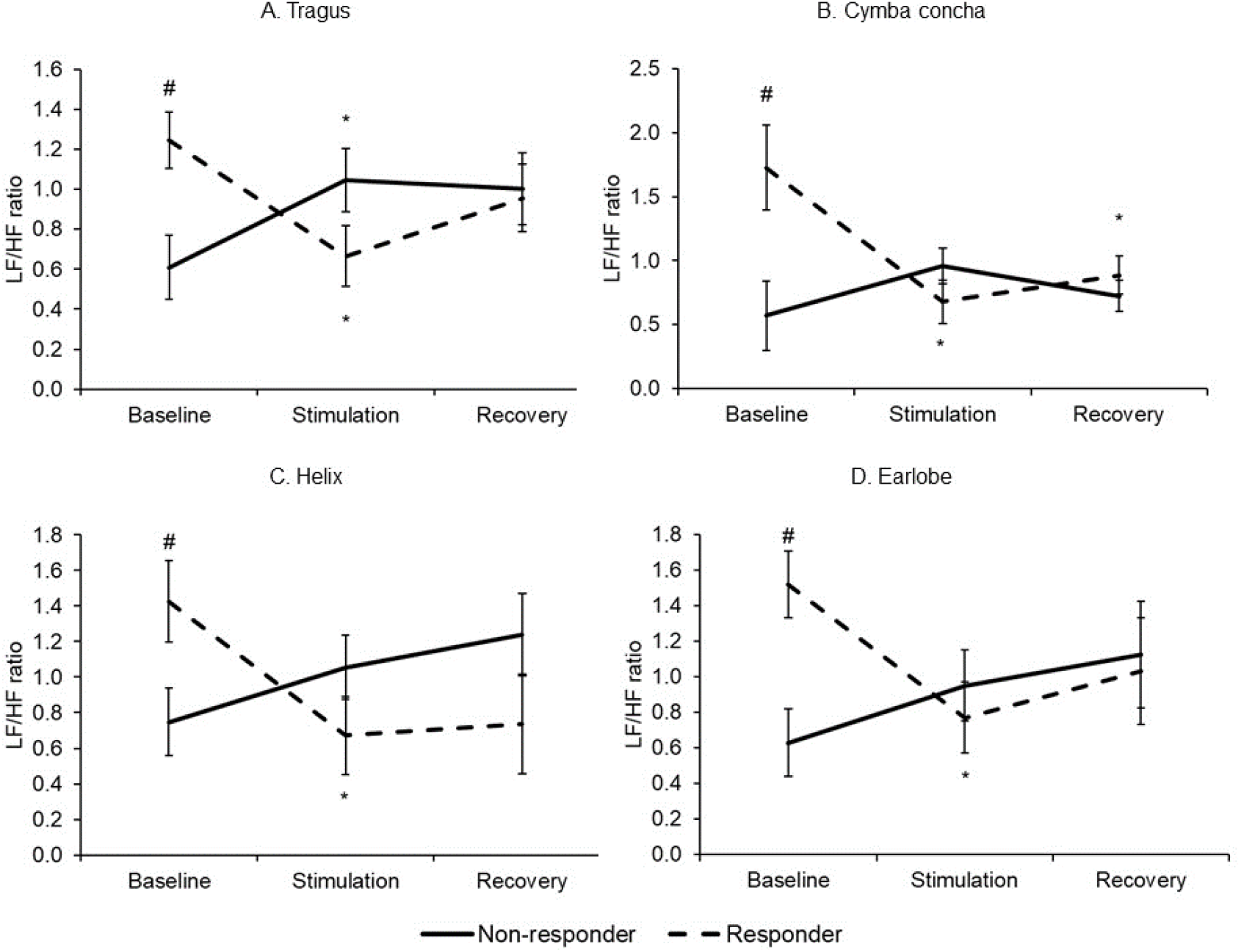
Significant interaction effects emerged for the LF/HF ratio when comparing response against conditions for stimulation at the tragus (A), cymba concha (B), helix (C) and earlobe (D). * = significantly different to baseline; # = significantly different to non-responder. Data presented as mean ± 1 SEM.

Baseline LF/HF ratio was significantly higher in responders than non-responders for stimulation administered to the tragus (p = 0.007), cymba concha (p = 0.014), helix (p = 0.035) and earlobe (p = 0.004). Furthermore, in the group of responders, LF/HF ratios significantly decreased during stimulation compared to baseline for tragus (p < 0.001), cymba concha (p = 0.005), helix (p = 0.001) and earlobe (p < 0.001). For tragus stimulation, there was a significant increase in LF/HF ratio between baseline and stimulation for the **group of non-responders** (p < 0.001). Furthermore, for stimulation delivered to the cymba concha, LF/HF ratio was significantly lower during recovery than baseline in the group of responders (p = 0.005).

### Reported sensations during stimulation

The electrical current was typically described at all stimulation sites as being a ‘tingling’, ‘tickling’, ‘pins and needles’ or ‘sharp’ sensation before the current was adjusted to a lower level. The sensations were also often localised to the stimulation site. However, some participants reported sensations at other sites on the ear or on nearby facial regions. For instance, the sensation evoked by helix stimulation was reported by some participants to travel posteriorly along the helix of the ear from the stimulation site (n = 5) or travel inferiorly towards the tragus on one or both ears (n = 3) or along the ipsilateral mandible (n = 2). Additionally, in two participants the sensation caused by earlobe stimulation was felt to radiate inferiorly along the neck, while one participant reported an itching sensation along their left zygomatic arch, which was not present on the right zygomatic arch. For tragus stimulation, three participants reported a tingling sensation, which radiated inferiorly along the mandible on one or both sides. Lastly, in two participants, cymba concha stimulation evoked a tingling or tickling sensation which radiated laterally from the cymba concha towards the helix of the ear and one participant reported a ‘vague sensation of current’ travelling inferiorly from the ear along the neck. One participant reported a sensation of ‘warmth’ in their chest for the duration of cymba concha stimulation, which ceased when the current was switched off.

## DISCUSSION

Transcutaneous auricular nerve stimulation (tANS) acutely administered for 15 minutes in healthy volunteers at the tragus, cymba concha, earlobe and helix significantly increased HRV measures. This agrees with our hypothesis that tragus, cymba concha and earlobe would change HRV but contrasts with our prediction that helix stimulation would be less effective than stimulation at the other sites. Interestingly, in line with our second hypothesis, linear regression modelling revealed that baseline LF/HF ratio significantly predicted change in LF/HF ratio (between baseline and stimulation) for all four ear sites and for the final sample regardless of site. This was characterised by a negative relationship where high LF/HF ratios at baseline were associated with greater decreases in LF/HF ratio with tANS. Consistent with these findings, response type (responder: decrease ≥ 20% in LF/HF ratio; non-responder: any increase or a < 20% decrease in LF/HF ratio) significantly interacted with condition suggesting that different responses between participants may be influenced by baseline values. Importantly, this reinforces the need to evaluate how HRV is modulated by tANS in individual participants (as well as the whole group), especially when considering the therapeutic applications of tANS.

### Effects of tANS on measures associated with autonomic function indicate a shift towards parasympathetic predominance

This study shows that tANS increases measures of HRV associated with parasympathetic predominance (2), such as time-domain HRV measures including RR interval, Δ RR interval, SDRR, RMSSD and pRR50, complemented by increases in non-linear indices such as (n)SD1 and SD2. This is in agreement with our previous studies showing that tANS applied to the tragus has been associated with increases in spontaneous BRS (6,9) while reductions in muscle sympathetic nerve activity are also observed (MSNA, (5)). Furthermore, tragus stimulation also increased Δ RR interval and improved HRV age in patients with tinnitus (8) and decreased LF/HF ratios in patients with diastolic dysfunction (7). There were some differences between the apparent extent of effect on frequency-domain HRV measures between this study and that of our previous work (5), although it is comparable to our more recent work investigating the effects of acute and chronic tANS in healthy older volunteers (9). Differences in the algorithms used to derive the frequency-domain HRV measures may underlie these discrepancies. For instance, the Fast Fourier Transform in Spike2 was used in Clancy et al. (2014) (5) whilst the Lomb-Scargle periodogram (used by default in LabChart8) was employed here and in our 2019 paper (9). Thus, discrepancies in the effects observed in prior work, may also be due to different HRV analysis programs and the different algorithms employed. To echo the methodological recommendations advocated by others (29), future research should therefore thoroughly report the software used to derive HRV including the algorithms used to derive frequency-domain HRV parameters.

### Is there an optimal site of electrode placement for influences on HRV by tANS?

This study explored how HRV measures compared when the same stimulation parameters were delivered to ear sites with presumed different innervation patterns. As the cymba concha, tragus, helix and earlobe are frequently used sites and thought to have overlapping innervation (ABVN: cymba concha and tragus; ATN: tragus and helix; GAN: tragus and earlobe), identical stimulation protocols were delivered to these four sites. We found no significant differences in responses between the four ear stimulation sites, with clear increases in HRV measures with tANS of all sites. This is an interesting finding that warrants further investigation to more fully understand the neural pathways involved in these effects of tANS.

As findings seem to suggest that there is no optimal site (with the stimulation parameters and measures employed here), one question is whether activation of the ABVN is **required** for these effects, as commonly thought. In an fMRI study, stimulation of the cymba concha (considered only vagally innervated) influenced activity in the nucleus of the tractus solitaries (NTS), a region of dense vagal afferent termination, while earlobe stimulation (potentially supplied by GAN afferents) activated different brain sites (32). Additionally, cymba concha stimulation produced stronger activation in the area of the medulla corresponding to the NTS compared to stimulation applied to the earlobe and inner tragus (presumed equally innervated by the ABVN and GAN) (33). However, in a rat study, where stimulation of the tragus elicited an inhibition of sympathetic nerve activity, neuronal tracing revealed that the majority of afferent labelling was not in fact localized in the NTS but instead could be identified in the cervical cord dorsal horn. Furthermore, the stimulation-evoked sympathoinhibition was abolished by sectioning the cervical sensory nerves (22), suggesting that the ABVN may not be necessary for changes in measures associated with autonomic function. Indeed, as suggested by differences between participants in the subjective responses to stimulation of the same site, innervation of the ABVN to the external auricular dermatome may be more complex due to this inter-individual variation (4,34). Of course, if the desired outcomes are achieved, such as improvements in autonomic function and/or therapeutic benefits, it could be considered unnecessary to know the exact nerves being stimulated.

### Bilateral vs unilateral auricular stimulation?

Few studies stimulate both left and right auricular sites simultaneously. Our finding that bilateral stimulation at any of the tested sites increased HRV is consistent, albeit not identical, to previous reports. For example, unilateral stimulation at different sites of the ear increased HRV (35) with greater effects of stimulation of the right ear compared to the left (23). Another study also reported a dominance of the right ear, since stimulation of the right but not left concha resulted in a significant increase in only one marker of HRV, SDNN, although left side stimulation had a tendency to increase HRV (16). Similarly, unilateral stimulation of either the concha or the tragus increased some HRV measures (36). It is tempting to speculate that simultaneous bilateral stimulation could provide more reliable effects on HRV than unilateral stimulation, but more research seems required before this can be concluded.

### Baseline HRV measures predict response to tANS

Although it is fundamental to evaluate how measures of autonomic function change with tANS for an entire sample of participants, it is also crucial to consider how responses to the stimulation differ between individuals. This is particularly important given the increasing interest in applying tANS in conditions that are characterised by impaired autonomic function. Indeed, previous work indicates that baseline LF/HF ratio was significantly associated with direction and magnitude of change with tANS delivered to the tragus (5,9). Consistent with this prior work, this study revealed that for each site and for all sites combined, baseline LF/HF ratio significantly predicted response with higher baseline LF/HF ratios being associated with greater decreases in LF/HF ratios during stimulation.

Combined, these findings suggest that starting values for measures of autonomic function may hold the key for determining the efficacy of tANS. Furthermore, starting measures reflecting autonomic function may provide the reason why some studies have seen little effect of tANS on measures associated with autonomic function: impaired values are less likely in young healthy volunteers compared to clinical samples, resulting in smaller (if any) effects.

Positive effects of tANS have been reported in conditions characterised by autonomic imbalance. For instance, in individuals with coronary artery disease, cymba concha stimulation for 15 minutes every day for two weeks improved exercise tolerance and reliance on vasodilator medication (37,38). In individuals with paroxysmal atrial fibrillation, tragal stimulation significantly reduced plasma levels of TNF-α, a pro-inflammatory cytokine, along with atrial fibrillation (10). tANS also appears to bring benefit in depression: stimulation applied for two weeks (39) and one month (40) in patients with depression significantly reduced depression scores. Therefore, further work is needed to explore the reproducibility of using a baseline measure of autonomic balance to predict the direction and magnitude of response in groups that experience imbalance in autonomic function.

Further support for the importance of considering how starting values of autonomic function influence change with tANS comes from comparing how response type to tANS interacted with the stimulation. Indeed, consistent with our previous studies (5,9), participants were classified as a responder if their LF/HF ratio decreased ≥ 20% between baseline and stimulation; or as a nonresponder if there was any increase or < 20% decrease in LF/HF ratio between baseline and stimulation. Findings revealed that responders had higher baseline LF/HF ratios than non-responders with values decreasing with stimulation in the former group. This therefore renders it unlikely that the observed effects were due to regression to the mean and/or participants lying down and experiencing a cumulative physiological relaxation effect. In addition, although the physiological interpretation of the LF/HF ratio is controversial, partly due to a lack of clarity concerning what LF HRV power really means (27–29), these findings are still noteworthy as they reinforce the importance of investigating why some individuals respond whilst others do not. Indeed, there is a focus in the literature on analysing the entire group, which may slow the application of tANS in conditions that are in need of safer therapies. By adopting a more individualised approach to tANS and undertaking further research exploring optimum stimulation parameters, it may be possible to further elucidate the mechanisms involved and bring health improvements to those who would benefit from tANS.

## Conclusion

This study revealed that stimulation administered to the tragus, cymba concha, helix or earlobe significantly influenced HRV measures towards parasympathetic prevalence regardless of ear site. Heightened LF/HF ratio at baseline was associated with greater decreases in LF/HF ratio with stimulation to all four sites, reinforcing the need for future work to further explore the influence of inter-individual variability in tANS. Findings also have implications for adopting a tailored approach when considering the therapeutic uses of tANS. A recent systematic review has highlighted some similar points (41) and in agreement with one major outcome of that review, we conclude that further research in optimising tANS stimulation parameters, the most effective set-up for specific target groups may be established.

## Acknowledgements

We would like to thank all of the volunteers who took part in the study.

