## Supplementary Table 1 for "The autonomic effects of transcutaneous auricular nerve stimulation at different sites on the external auricle of the ear"

Supplementary Table 1: Summary of all main effects and interaction effects.

| | | Main effect of condition | Main effect of ear site | Condition*ear site interaction |
|---|---|---|---|---|
| Time HRV | RR interval | $F(1.16,24.37) = 33.66$, $p < 0.001$, $\eta_p^2 = 0.616$ | $F(1.98,41.47) = 0.91$, $p > 0.05$, $\eta_p^2 = 0.042$ | $F(3.51,73.68) = 2.04$, $p > 0.05$, $\eta_p^2 = 0.089$ |
| | Δ RR | $F(2,42) = 10.96$, $p < 0.001$, $\eta_p^2 = 0.343$ | $F(3,63) = 2.01$, $p > 0.05$, $\eta_p^2 = 0.087$ | $F(4.07,85.51) = 1.02$, $p > 0.05$, $\eta_p^2 = 0.046$ |
| | SDRR | $F(1.29,27.07) = 15.17$, $p < 0.001$, $\eta_p^2 = 0.419$ | $F(3,63) = 2.05$, $p > 0.05$, $\eta_p^2 = 0.089$ | $F(3.71,77.86) = 0.83$, $p > 0.05$, $\eta_p^2 = 0.038$ |
| | RMSSD | $F(1.16,24.29) = 12.11$, $p = 0.001$, $\eta_p^2 = 0.366$ | $F(1.88,39.43) = 1.41$, $p > 0.05$, $\eta_p^2 = 0.063$ | $F(3.18,66.81) = 0.64$, $p > 0.05$, $\eta_p^2 = 0.030$ |
| | pRR50 | $F(1.44,30.26) = 11.36$, $p = 0.001$, $\eta_p^2 = 0.351$ | $F(2.25,47.18) = 0.57$, $p > 0.05$, $\eta_p^2 = 0.026$ | $F(4.06,85.16) = 1.31$, $p > 0.05$, $\eta_p^2 = 0.059$ |
| Freq HRV | Total power | $F(1.32,27.68) = 5.62$, $p = 0.007$, $\eta_p^2 = 0.211$ | $F(1.97,41.42) = 1.88$, $p > 0.05$, $\eta_p^2 = 0.082$ | $F(3.91,82.11) = 0.82$, $p > 0.05$, $\eta_p^2 = 0.038$ |
| | LF power | $F(1.22,25.61) = 4.58$, $p = 0.035$, $\eta_p^2 = 0.179$ | $F(2.11,44.38) = 1.52$, $p > 0.05$, $\eta_p^2 = 0.068$ | $F(2.50,52.57) = 0.65$, $p > 0.05$, $\eta_p^2 = 0.030$ |
| | HF power | $F(1.21,25.37) = 4.92$, $p = 0.030$, $\eta_p^2 = 0.190$ | $F(1.98,41.52) = 1.58$, $p > 0.05$, $\eta_p^2 = 0.070$ | $F(1.44,30.17) = 0.59$, $p > 0.05$, $\eta_p^2 = 0.027$ |
| | LF/HF ratio | $F(2,42) = 1.24$, $p > 0.05$, $\eta_p^2 = 0.056$ | $F(3,63) = 0.37$, $p > 0.05$, $\eta_p^2 = 0.018$ | $F(3.33,69.94) = 0.57$, $p > 0.05$, $\eta_p^2 = 0.027$ |
| Non-linear HRV | S | $F(1.07,22.46) = 7.49$, $p = 0.011$, $\eta_p^2 = 0.263$ | $F(1.56,32.76) = 1.38$, $p > 0.05$, $\eta_p^2 = 0.062$ | $F(3.09,64.79) = 0.89$, $p > 0.05$, $\eta_p^2 = 0.041$ |
| | SD1 | $F(1.16,24.29) = 12.12$, $p = 0.001$, $\eta_p^2 = 0.366$ | $F(1.88,39.41) = 1.42$, $p > 0.05$, $\eta_p^2 = 0.063$ | $F(3.18,66.83) = 0.64$, $p > 0.05$, $\eta_p^2 = 0.030$ |
| | nSD1 | $F(1.20,25.09) = 9.87$, $p = 0.003$, $\eta_p^2 = 0.320$ | $F(2.11,44.32) = 1.35$, $p > 0.05$, $\eta_p^2 = 0.060$ | $F(3.41,71.60) = 0.47$, $p > 0.05$, $\eta_p^2 = 0.022$ |
| | SD2 | $F(1.39,29.21) = 14.72$, $p < 0.001$, $\eta_p^2 = 0.412$ | $F(3,63) = 2.15$, $p > 0.05$, $\eta_p^2 = 0.093$ | $F(3.64,76.43) = 0.90$, $p > 0.05$, $\eta_p^2 = 0.041$ |
| | nSD2 | $F(2,42) = 10.76$, $p < 0.001$, $\eta_p^2 = 0.339$ | $F(3,63) = 2.34$, $p > 0.05$, $\eta_p^2 = 0.100$ | $F(3.70,77.77) = 0.84$, $p > 0.05$, $\eta_p^2 = 0.038$ |
| BRS | Up BRS | $F(2,34) = 3.06$, $p = 0.060$, $\eta_p^2 = 0.153$ | $F(3,51) = 2.26$, $p > 0.05$, $\eta_p^2 = 0.117$ | $F(2.24,38.11) = 0.94$, $p > 0.05$, $\eta_p^2 = 0.053$ |
| | Down BRS | $F(2,34) = 0.74$, $p > 0.05$, $\eta_p^2 = 0.042$ | $F(3,51) = 1.15$, $p > 0.05$, $\eta_p^2 = 0.063$ | $F(3.36,57.18) = 1.43$, $p > 0.05$, $\eta_p^2 = 0.077$ |
| | Mean BRS | $F(1.48,25.17) = 3.34$, $p = 0.065$, $\eta_p^2 = 0.164$ | $F(2.02,34.35) = 1.26$, $p > 0.05$, $\eta_p^2 = 0.069$ | $F(2.82,47.90) = 0.97$, $p > 0.05$, $\eta_p^2 = 0.054$ |
