## Supplementary Table 2 for "The autonomic effects of transcutaneous auricular nerve stimulation at different sites on the external auricle of the ear"

Supplementary Table 2: Summary of linear regressions for each ear site with baseline values as the independent variable and change (Δ, between baseline and stimulation) as the dependent variable.

| | | Tragus | | Earlobe | | Helix | | Cymba concha | |
|---|---|---|---|---|---|---|---|---|---|
| | | $R^2$ | p-value | $R^2$ | p-value | $R^2$ | p-value | $R^2$ | p-value |
| **Time HRV** | **RR interval** | 0.000 | > 0.05 | 0.012 | > 0.05 | 0.154 | 0.071 | 0.320 | 0.006 |
| | **Δ RR** | 0.011 | > 0.05 | 0.000 | > 0.05 | 0.032 | > 0.05 | 0.003 | > 0.05 |
| | **SDRR** | 0.021 | > 0.05 | 0.004 | > 0.05 | 0.223 | 0.027 | 0.016 | > 0.05 |
| | **RMSSD** | 0.000 | > 0.05 | 0.173 | 0.054 | 0.008 | > 0.05 | 0.125 | > 0.05 |
| | **pRR50** | 0.000 | > 0.05 | 0.026 | > 0.05 | 0.003 | > 0.05 | 0.137 | > 0.05 |
| **Freq HRV** | **Total power** | 0.121 | > 0.05 | 0.052 | > 0.05 | 0.421 | 0.001 | 0.023 | > 0.05 |
| | **LF power** | 0.051 | > 0.05 | 0.068 | > 0.05 | 0.611 | < 0.001 | 0.541 | < 0.001 |
| | **HF power** | 0.270 | 0.013 | 0.284 | 0.011 | 0.155 | 0.070 | 0.000 | > 0.05 |
| | **LF/HF ratio** | 0.336 | 0.005 | 0.388 | 0.002 | 0.349 | 0.004 | 0.793 | < 0.001 |
| **Non-linear HRV** | **S** | 0.04 | > 0.05 | 0.236 | 0.022 | 0.177 | 0.051 | 0.485 | < 0.001 |
| | **SD1** | 0.000 | > 0.05 | 0.174 | 0.054 | 0.007 | > 0.05 | 0.126 | > 0.05 |
| | **nSD1** | 0.019 | > 0.05 | 0.126 | > 0.05 | 0.066 | > 0.05 | 0.021 | > 0.05 |
| | **SD2** | 0.027 | > 0.05 | 0.010 | > 0.05 | 0.316 | 0.006 | 0.022 | > 0.05 |
| | **nSD2** | 0.006 | > 0.05 | 0.097 | > 0.05 | 0.377 | 0.002 | 0.017 | > 0.05 |
| **BRS** | **Up BRS** | 0.168 | 0.058 | 0.037 | > 0.05 | 0.001 | > 0.05 | 0.077 | > 0.05 |
| | **Down BRS** | 0.021 | > 0.05 | 0.039 | > 0.05 | 0.050 | > 0.05 | 0.499 | < 0.001 |
| | **Mean BRS** | 0.057 | > 0.05 | 0.016 | > 0.05 | 0.003 | > 0.05 | 0.295 | 0.009 |
